# Linking microbial communities to ecosystem functions: what we can learn from genotype-phenotype mapping in organisms

**DOI:** 10.1101/740373

**Authors:** Andrew H. Morris, Kyle M. Meyer, Brendan J. M. Bohannan

## Abstract

Microorganisms mediate many important ecosystem functions, yet it remains unclear to what extent microbial diversity or community composition is important for determining the rates of ecosystem-scale functions. This uncertainty limits our ability to predict and manage crucial microbially-mediated processes, such as nutrient loss and greenhouse gas emissions. Our lack of understanding stems from the relatively large diversity of microorganisms, the difficulty in directly identifying functional groups, and our limited ability to manipulate microbial community attributes. For this reason, we propose that integrating traditional biodiversity-ecosystem function research with ideas from genotype-phenotype mapping could provide the new perspective our discipline needs. We identify three insights from genotype-phenotype mapping that could be useful for microbial biodiversity-ecosystem function studies: the concept of “agnostic” mapping, the use of more powerful ways to account for multiple comparisons, and the incorporation of covariates into models of ecosystem function. We illustrate the potential for these approaches to elucidate microbial biodiversity-ecosystem function relationships by analyzing a subset of published data measuring methane oxidation rates from incubations of tropical soil. We assert that combining the approaches of traditional biodiversity-ecosystem function research with ideas from genotype-phenotype mapping will not only generate novel hypotheses about how complex microbial communities drive ecosystem function, but also help scientists predict and manage changes to ecosystem functions resulting from human activities.

## 1 Introduction

### 1.1 A new perspective on microbial community-ecosystem function relationships is needed

Ecologists have long investigated the effects of changing biodiversity on ecosystem function, documenting, for example, relationships between terrestrial plant community richness and primary productivity (1). The study of the relationship between microbial biodiversity and ecosystem functions is much more recent and has generated mixed results (2,3). Despite the fact that microbes mediate many important ecosystem functions, it remains maddeningly unclear to what extent microbial diversity or community composition is important for determining the rates of ecosystem-scale functions. For example, numerous studies have attempted to correlate microbial functional group abundance or diversity with the rate of various ecosystem functions and most are unsuccessful (4,5). This uncertainty limits our ability to predict and manage crucial microbially-mediated ecosystem functions, such as nutrient loss and greenhouse gas emissions. As a discipline, we need a new perspective on these questions.

Our lack of understanding stems from many sources. Microbial communities are much more diverse than those of plants and more abundant, and thus more difficult to sample comprehensively. Unlike plants, functional groups of microbes are rarely determined through observation or direct measurement, but rather must be indirectly inferred from environmental DNA, adding uncertainty. The ability to experimentally manipulate microbial diversity or community composition is much more limited for microbes. Given all of these differences (and many others), the challenge of linking microbial communities to ecosystem functions may be less like that of plant communities, and more analogous to the complex task of linking genomic variation to organismal phenotypes. For this reason, we propose that integrating traditional biodiversity-ecosystem function research with ideas from genotype-phenotype mapping could provide the new perspective our discipline needs.

### 1.2 There is evidence that microbial diversity can matter

Perhaps our uncertainty arises because there is no general relationship between microbial biodiversity and any particular ecosystem function. For example, if local selection always optimizes the available microbial biodiversity to maximize ecosystem function, then microbial community composition should not matter for predicting the rates of microbially-mediated functions. Instead, the rates of these functions should be determined by the underlying environmental variation. In this case, the microbial community would simply act as a conduit through which the abiotic environment alters ecosystem function. Attempts to document a general relationship between microbial community attributes and function would then fail (since the primary drivers are environmental).

However, there is evidence that this is not always the case, A limited number of studies have manipulated the connection between environmental factors and microbial community composition through “common garden” or reciprocal transplant experiments, and they frequently report different rates of ecosystem functions for different microbial communities under the same environmental conditions (6). This has been observed for plant decomposition (7,8), plant phenology (9), soil nitrogen cycling (10), and soil greenhouse gas emissions (11), among others. Therefore, variation in microbial community composition can be associated with variation in ecosystem function, independent of environmental variation. So why has this been difficult to consistently document in the field?

### 1.3 Traditional approaches to quantifying this relationship have provided only minor improvements

Most comparative studies of microbial community function in the field focus on one of two aspects of microbial community structure that are hypothesized to predict ecosystem function. The first aspect is “functional” gene or transcript abundance. In this case, qPCR or shotgun metagenomic sequencing is used to estimate the abundance of a gene or transcript that is a putative marker for a microbial process (and thus a marker for the functional group that performs this process). For example, the gene *mcrA*, which encodes a subunit of the enzyme that performs the final step in methanogenesis, is commonly used as a marker for methanogenesis and for the methanogen functional group. Other examples include *pmoA* and methanotrophy, *nifH* and nitrification, and *nosZ* and denitrification. The abundance of these markers is often hypothesized to be predictive of the rate of their associated processes (for example, the abundance of *mcrA* is often hypothesized to be related to the rate of methanogenesis). There are examples where this relationship is present (12–14). However, a review of such studies found that the abundance of a functional gene or transcript is rarely correlated with the rate of the corresponding process and depends on the function of interest, with most effects either negative or not significant (4).

The second aspect of microbial community structure hypothesized to predict ecosystem function is taxonomic or functional diversity. Diversity is either estimated from sequence variants of a barcode gene such as the 16S rRNA gene or manipulated through some proxy of diversity such as sequential dilution or varying filter sizes. Studies that experimentally manipulate diversity generally find a relationship between the applied diversity treatment and ecosystem functions, including for methanotrophy (14), phosphorus leaching (15), greenhouse gas emissions (15), decomposition (16), and nitrogen cycling (17). However, in many cases, richness is confounded with other factors such as the presence or absence of major phylogenetic groups (nematodes, fungi) or abundance of microbial cells. In addition, assembled microbial communities pose the same challenges as macroorganismal diversity experiments, for example the highest diversity treatment will more likely contain the most productive taxon. A review of microbial diversity studies shows that microbial community taxonomic and functional diversity add little explanatory power to models of ecosystem function (5). Overall, functional gene abundance and community diversity improve models of ecosystem function less than one third of the time and increase variance explained by an average of only 8 percentage points (5).

### 1.4 Genotype-phenotype mapping as a source of inspiration

Given the relative lack of success to date, it is time to rethink how we approach the challenge of mapping microbial community structure to ecosystem function. Microbial ecologists are not the only biologists who are attempting to determine the relationship between a complex set of highly-variable data and an aggregate function. This kind of “many-to-one” mapping is analogous to the challenge of identifying the genetic basis of complex traits in organismal populations. In such “genotype-phenotype” mapping studies, a population exhibits variation in a phenotype (e.g. height or disease state) as well as variation in potentially thousands of single nucleotide polymorphisms (SNPs). To identify the genetic basis for a trait, investigators sample from this population and correlate phenotype with genotype. In most cases, there are many more loci than individuals and we do not know whether the SNPs are causally linked or are simply in linkage disequilibrium with a causal mutation.

There are a number of parallels between this challenge faced by organismal biologists and that facing microbial community ecologists. They are both “many-to-one” mapping challenges, which involve large numbers of statistical comparisons. Both are attempting to identify causal relationships that are potentially confounded by very complex patterns of covariation. There is often no strong expectation about which entities (i.e. which genomic regions or which microbial taxa) are most likely to be causally-related to phenotype or function, and thus “agnostic” approaches are needed. Both are sensitive to exactly how the “mapping” question is asked, and ultimately require manipulation (of genes or taxa) to establish causation. We describe each of these parallels below in more detail, and provide an example of how these ideas could be applied to microbial data.

### 1.5 The importance of a taxonomically “agnostic” approach

The traditional approach to microorganismal biodiversity-ecosystem function research is to measure or manipulate the diversity of a taxonomic group (e.g. plants) and look for an association with the function performed by that group (e.g. primary productivity). In the broadest sense, we can think of plants as a “functional group”, i.e. a group of taxa united by their ability to perform a particular ecosystem function. Ecologists may further divide a functional group (e.g. plants) into smaller functional groupings (e.g. forbs), defined by the details of how they perform their particular ecosystem function.

However, for microbes our knowledge of functional groups is much more limited. From a very limited number of cultured isolates we have a provisional understanding of which microbes might be involved in some ecosystem functions. And by sequencing the genomes of these isolates, we have identified genetic markers for certain functions. But most microbial taxa remain uncultured and we do not know the function of most microbial taxa detected in environmental samples (18,19). In addition, there have been recent discoveries of functions in unexpected taxonomic groups, for example methanogenesis by fungi and cyanobacteria, a function previously considered restricted to the Archaea (20,21). Because of this, it would be prudent to look more agnostically at microbial communities to identify taxa or groups of taxa that are important for predicting the rate of ecosystem functions rather than assuming that the genetic markers we have provisionally identified for a given function represent the most likely taxa involved. This agnostic approach is analogous to the approach of many genotype-phenotype mapping studies (e.g. genome-wide association studies, aka GWAS), which often look for associations between a phenotype and loci anywhere in a genome.

### 1.6 Dealing with multiple comparisons and covariation

Microbial biodiversity-ecosystem function studies and genotype-phenotype mapping studies are both “many-to-one” mapping challenges, which involve large numbers of statistical tests. Hundreds to thousands of tests are routinely made per study, greatly inflating the number of false positives identified using statistical hypothesis testing approaches. Some microbial biodiversity-ecosystem function studies do not correct for this, while others use approaches that may unnecessarily inflate the false negative rate, such as the Bonferroni correction. The Bonferroni correction has been widely considered to be too conservative, particular for exploratory studies designed to generate hypotheses (22–24). Statisticians have developed a number of less-conservative approaches to correct for multiple comparisons by controlling the false discovery rate in order to balance the tradeoff between Type I and Type II errors (25,26). These approaches are commonly used in genotype-phenotype mapping studies.

It is widely accepted that organisms, including microorganisms, exhibit population stratification due to geographic and environmental separation (27,28). Genome-wide association studies generally account for population structure due to shared ancestry among cases and controls when modeling the connection between genotype and phenotype. The classic example is the latitudinal gradient of both height and genotypic similarity in Europe, which results in spurious associations between human height and genetic variation (29,30). To correct for this covariance structure, GWAS models incorporate genotypic similarity to correct for shared ancestry using a variety of methods, such as principal component correction or variance component modeling (31,32). Typically, microbial biodiversity-ecosystem function studies do not account for population stratification (i.e. community similarity across samples), although there are some exceptions (33,34). GWAS generally ignores the underlying environmental and spatial distance between samples and instead uses shared ancestry as a proxy for these variables. However, community similarity (the community analogue of shared ancestry among organisms) is not as tightly linked to geography or environment as is shared ancestry among organisms, and it could be very useful to account for these separately in microbial studies, especially if one is particularly interested in how composition alters function independent of the underlying environmental variation.

## 2 An example: high-affinity methane oxidation

To illustrate the ideas outlined above, we reanalyzed a subset previously published data (33) using a modified procedure from the original version. A full description of the study design, samples, and data generation can be found in that manuscript. Briefly, these data were gathered from intact soil cores taken from diverse ecosystems of the Congo Basin in Gabon, Africa. Cores were incubated in the laboratory under different concentrations of methane to identify the rates of specific methane cycling pathways. For this example, we will analyze data from just one of these pathways, high-affinity methane oxidation (the oxidation of atmospheric concentrations of methane), which we will refer to below as “methane oxidation”. In addition, for simplicity we only include amplicon sequences from the DNA-inferred community and not the RNA-inferred community, both of which are presented in the original paper (33). The data we analyzed include methane oxidation measurements, amplicon sequence variants (ASVs; (35)) generated using *DADA2* and inferred from 16S rRNA gene sequences, *pmoA* abundance estimates (via qPCR), latitude and longitude, and four environmental covariates (soil moisture, bulk density, carbon, and nitrogen).

Analyses were conducted in the *R* statistical environment using the *phyloseq* package (36,37). The relative abundances of ASVs were corrected using the variance stabilizing transformation from *DESeq2* (38,39). We first test typical measures of microbial structure including functional gene abundance and community richness, which was estimated using the *breakaway* package (40). We then demonstrate significant covariation between community structure (estimated as Bray-Curtis distance using *vegan*), environmental variation (euclidean distance), and geographic distance (euclidean distance) using Mantel tests (41,42). Finally, we present one approach to identifying taxa which are significantly associated with function independent of the environment by fitting variance component models using *varComp* to test the relationship between relative abundance of each ASV and methane oxidation rate (43). To illustrate how including different covariates (environmental, geographic, and community) can result in different conclusions about which taxa are associated with function, we fit this model with and without random effects variance components for environmental similarity, geographic site ID, and Bray-Curtis similarity. Significant taxa were determined by controlling the false discovery rate at q-value < 0.05 (26). Figures were created using *ggplot2* (44).

Methane oxidation rate was not significantly correlated with *pmoA* gene abundance or richness (Table 1, Figure 1). We tested collinearity between each pair of distance matrices for community, environment, and geography using Mantel tests and estimated p-values by permutation. We found a moderate and significant correlation between community composition and environmental variation, geography and community composition, and geography and environmental variation (Table 2, Figure 2). Principal coordinate plots show that beta diversity of samples separated by site ID and by ecosystem type (wetland or upland; Figure 2). Finally, we tested the effect of the relative abundance of each ASV on methane oxidation rate. We found different numbers of taxa significantly associated with methane oxidation depending on which covariates were included in the model (Table 3). In particular, 460 taxa were identified with no covariates whereas 6 were identified after including all covariates. Though we cannot infer function from 16S sequences, these 6 taxa fall into three genera and one class with cultured representatives that are not known to consume methane (45–48). Their effect on methane oxidation rate ranges from 0.5 to 1.5 (Figure 3). An effect of 1.5 means that for a one unit increase in relative abundance there is a 1.5 increase in k (the rate of exponential decrease in methane concentration over time).

**Table 1.** Functional gene abundance and ASV richness are not good predictors of methane oxidation rate. Linear models predicting methane oxidation rate from measures of microbial community structure.

| Term | Estimate | SE | t-statistic | p-value | n |
| --- | --- | --- | --- | --- | --- |
| *pmoA* copy number | 0.019 | 0.011 | 1.700 | 0.096 | 42 |
| Richness | 0.001 | 0.001 | 0.694 | 0.491 | 44 |

**Table 2.**
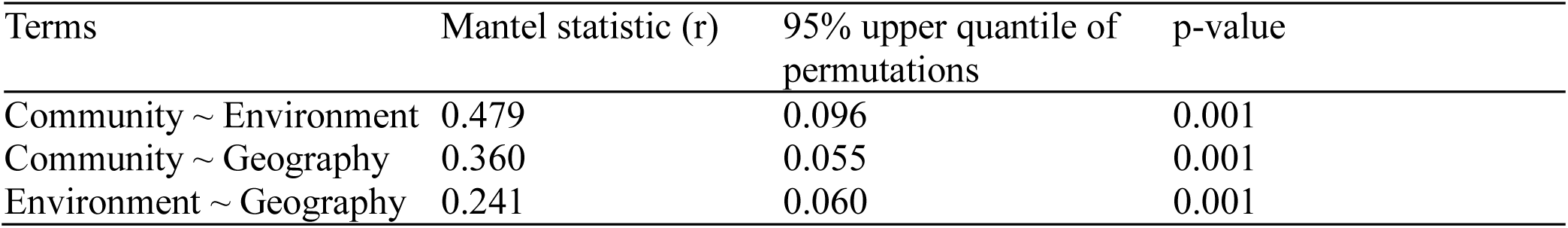
Mantel tests for each pair of dissimilarity matrices. Community distance matrix was based on Bray-Curtis distance while both environment and geography distance matrices were based on Euclidean distance. P-values determined by permutation test with 999 permutations.

**Table 3.**
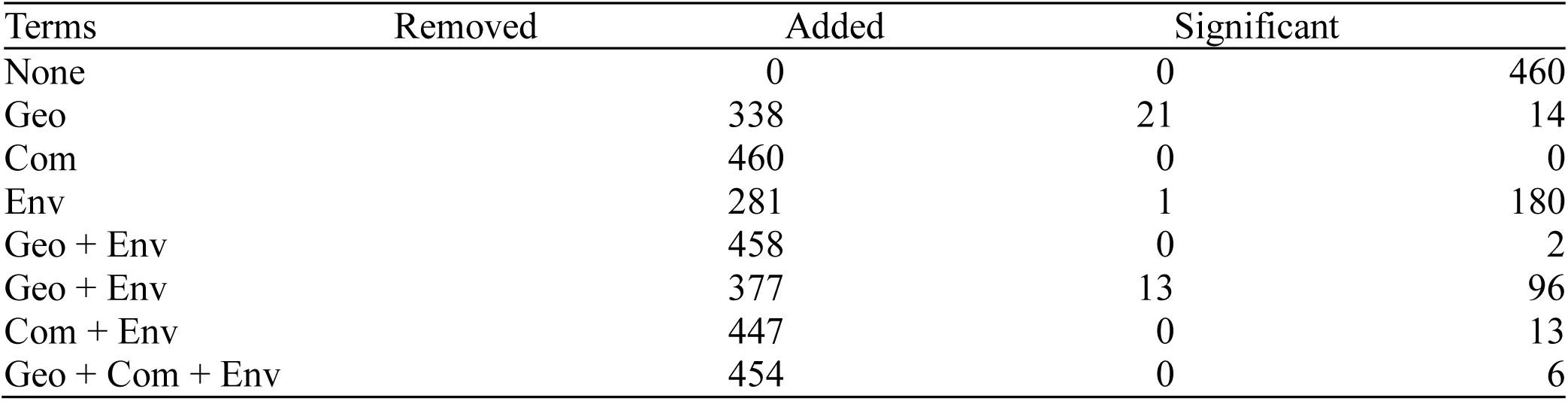
Number of significant taxa after including each set of covariates in a variance component model. Removed and Added columns are relative to the no-covariate model. Significance is determined by controlling the false discovery rate at q-values < 0.05.

**Figure 1.**
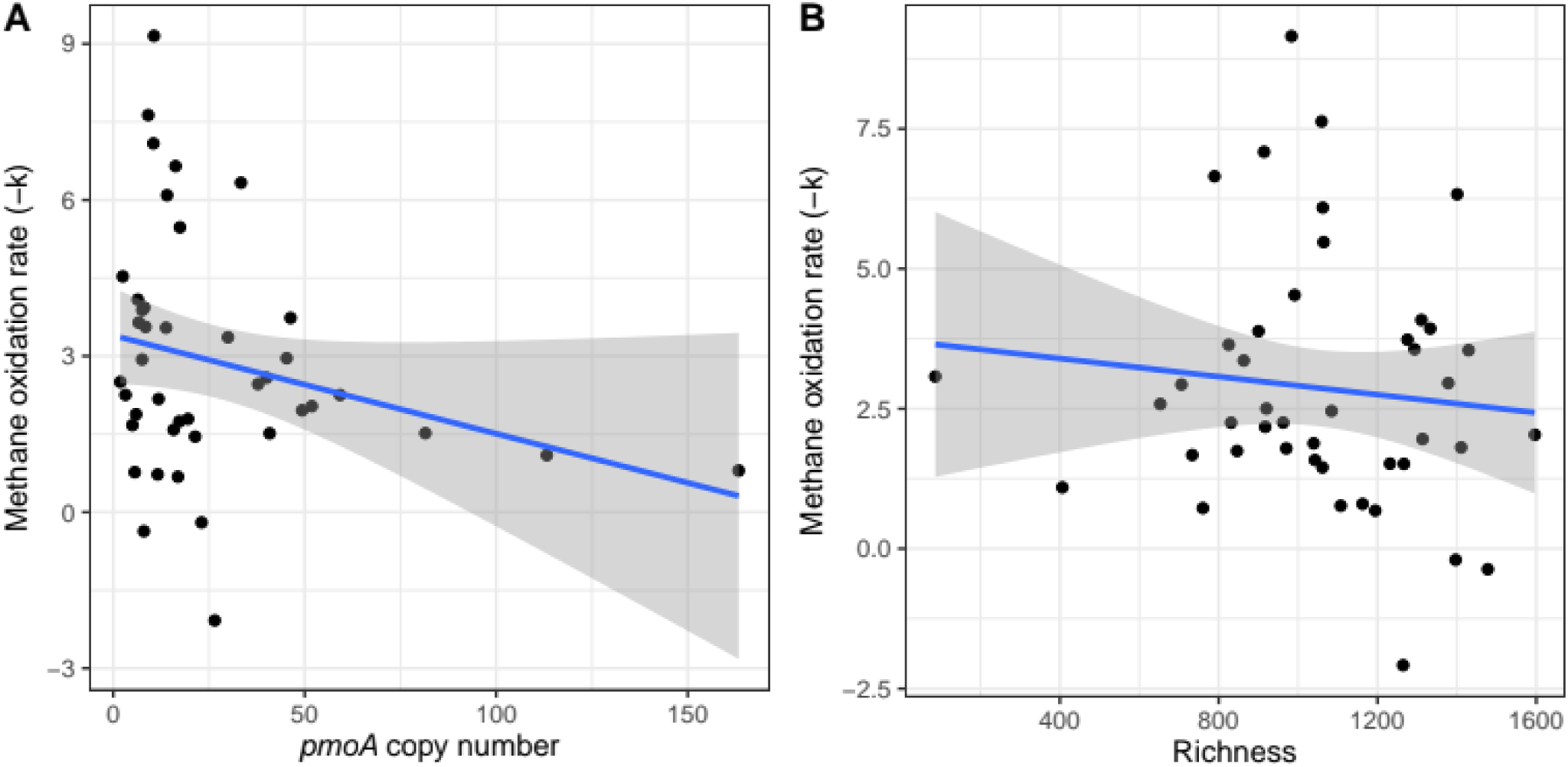
Methane oxidation rate is not correlated with functional gene abundance or ASV richness. Correlations between community attributes and ecosystem function. A) Abundance of the functional gene *pmoA* (n = 42) and B) ASV richness (n = 44). Lines represent the ordinary least squares regression line with standard errors.

**Figure 2.**
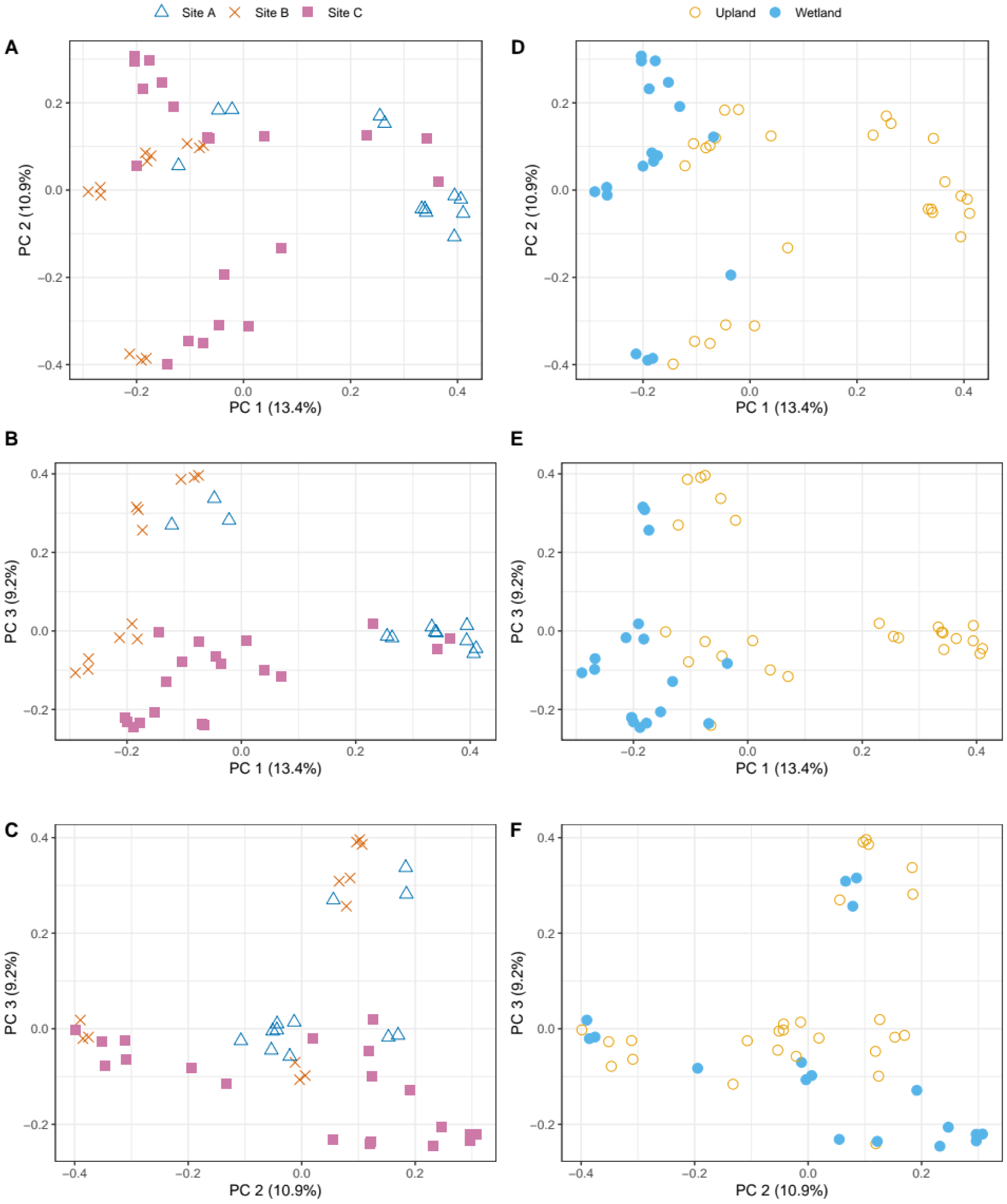
Microbial community composition is spatially and environmentally structured. Principle coordinate plots of Bray-Curtis distance representing the first three axes of community composition. In A, B, and C, points are identified by Site ID and in D, E, and F, points are identified by wetland or upland ecosystem. All four environmental covariates separate strongly by wetland/upland. Axis length is proportional to variance explained as indicated in parentheses. PC = principal coordinate.

**Figure 3.**
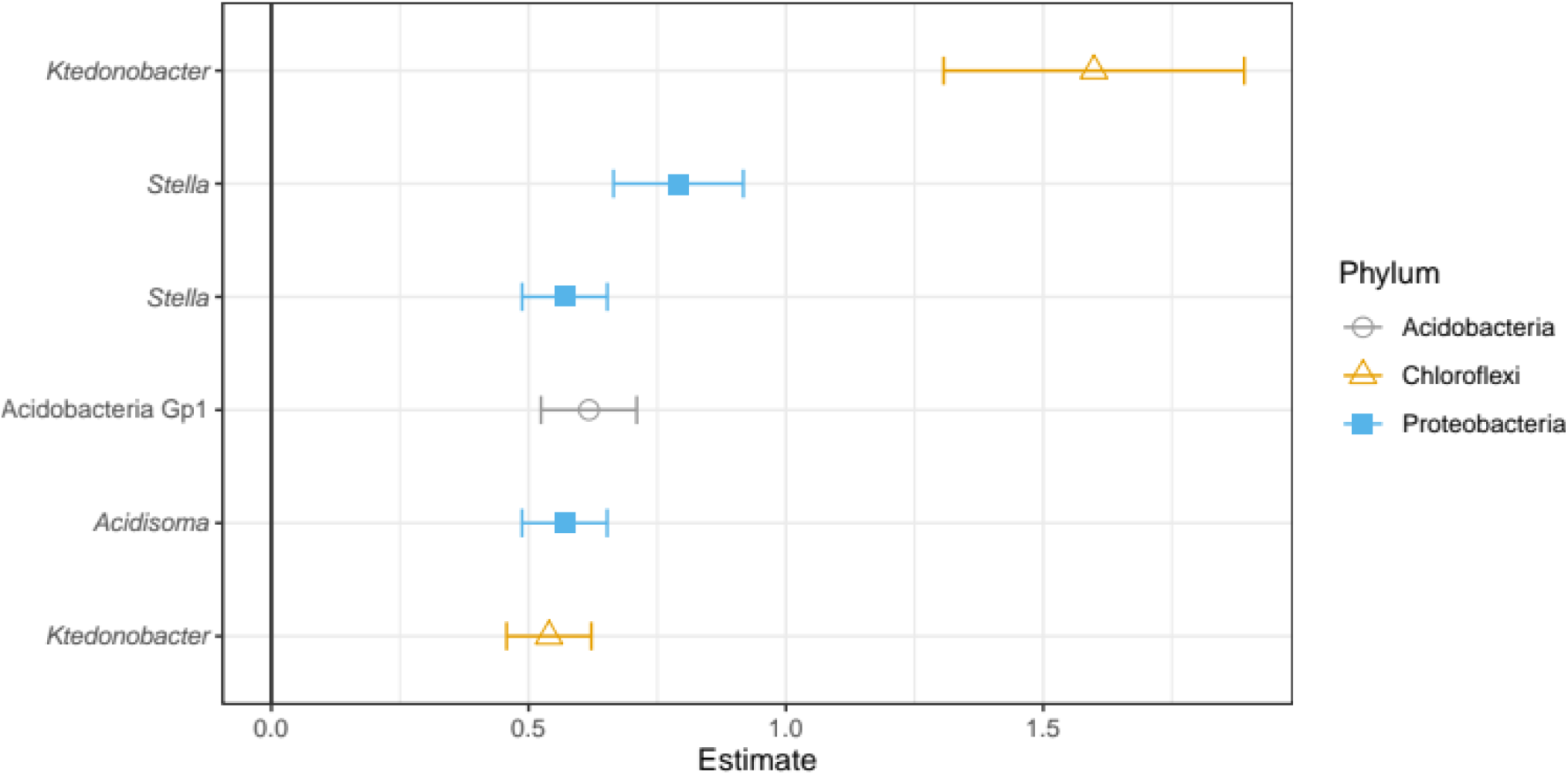
Taxa associated with methane oxidation rates after controlling for geographic location, environmental similarity, and community composition. Points are estimates for the linear relationship between the relative abundance of a single ASV and methane oxidation rate with standard errors from variance component models including similarity matrices as covariates for community and environment and site ID for geographic location. Amplicon sequence variants are labelled at the finest resolution available: genus for all except the Group 1 Acidobacterium. Points are identified by Phylum. Significant taxa were determined by controlling the false discovery rate at q-value < 0.05.

## 3 Discussion

In this paper, we argue that the traditional approach to microbial biodiversity-ecosystem function research often ignores the complexity of microbial community dynamics, in particular the complex patterns of covariation that often arise among microbial community similarity, environmental similarity, and spatial proximity. We propose that a better approach would be to learn from other complex many-to-one mapping problems in biology, particularly genotype-phenotype mapping. These studies often explicitly account for covariation, they correct for multiple comparisons in powerful ways, and they frequently take an “agnostic approach” that does not assume a particular relationship between structure and function.

We illustrate the potential for these approaches to elucidate microbial biodiversity-ecosystem function relationships by analyzing a subset of published data from incubations of tropical soil. In this example, soil cores from different ecosystems exhibited different rates of methane oxidation, but simple correlations between these rates and the diversity or abundance of putative methane oxidizing bacteria were not informative. These results agree with recent reviews of the literature that demonstrate that such simple approaches are often not a fruitful avenue for elucidating microbial structure-function connections.

We then asked if the relative abundance of particular taxa were related to ecosystem function, and we identified 460 taxa whose relative abundances were associated with rates of methane oxidation after controlling the false discovery rate (q-value < 0.05). These taxa could be related to ecosystem function in multiple ways. The most interesting possibility is that each of these taxa is statistically related because it is causally connected to the function. This could be direct, for example an organism that consumes methane, or indirect, for example an organism that regulates substrates necessary for consumers of methane. In either of these cases, the taxon could be useful as a biomarker of function or as an organism to investigate in order to better understand the biological drivers of variation in methane oxidation.

Alternatively, a significant association could occur for non-causal reasons. For example, any organism that tends to be in high abundance where methane oxidation rates are high would be correlated with methane oxidation, even if it has no causal relationship. This could be because such an organism is favored under the same environmental conditions that favor methane oxidizing bacteria (or that favor methane oxidation in general). Such covariation can drive associations that are not causal, but the effects of such covariation can be reduced through a number of approaches (many developed by biologists interested in genotype-phenotype mapping).

In our example, we showed that the abundances of microbial taxa exhibit complex patterns of covariation with each other and with environmental conditions and spatial location. These biogeographic patterns might be even stronger if we could sample the populations more intensively (49). Once we account for this covariation, our list of taxa associated with function was reduced to 6. These 6 taxa include taxa that are not known to directly contribute to methane oxidation, suggesting that the drivers of methane oxidation may be indirect, perhaps mediated by ecological interactions among taxa from multiple functional groups.

This approach is powerful because it accounts for the covariances that arise from the fundamental ecological processes that drive community assembly and underlie biogeographical patterns (27,50). It is also powerful because it is focused on a much more specific version of the “biodiversity-ecosystem function” question, i.e. it asks “which taxa are *uniquely* associated with function?”. By “uniquely associated”, we mean those taxa associated with function irrespective of environmental conditions, local community structure, or spatial proximity. This is not only a more specific question than is usually asked in microbial biodiversity-ecosystem function studies, but it is also a more appropriate one, especially if one is interested in how to incorporate microbial community data into ecosystem models. For modeling what is usually important is to identify those taxa that add explanatory power beyond that provided by other factors (such as environmental conditions).

We identified three insights from genotype-phenotype mapping that could be useful for microbial biodiversity-ecosystem function studies: the concept of “agnostic” mapping, the use of more powerful ways to account for multiple comparisons, and the incorporation of covariates. But there are other insights to be gained as well. For example, some phenotypes (e.g. the propensity for diseases such as Parkinson’s) are controlled by a single genetic locus (51,52). However, most phenotypes studied to date arise from the influence of many loci of small effect (as well as environmental factors; (53,54)). Organismal biologists have developed approaches tailored to identifying such loci; for example, by sampling organismal populations in a way that constrains genetic variation unrelated to phenotype. Similarly, for some ecosystem functions, it is possible that a single microbial taxon could substantially influence its rate. For example, methane flux from permafrost in Sweden may be controlled by a single taxon (55). However, most microbially-mediated ecosystem functions are likely the result of interactions (direct and indirect) among many taxa. We could maximize our ability to identify taxa of small effect by constraining variation, e.g. by sampling microbial communities in a way that makes them similar in structure (similar ecosystem, soil type, and abiotic conditions) while still varying in function.

Ultimately, the relationships identified in genotype-phenotype mapping studies must be verified. There are multiple ways that this verification is accomplished. In some cases, organisms can be artificially selected for a particular phenotype (e.g. through experimental evolution in an environment that favors the phenotype of interest) and the genetic changes that occur in response to selection can be compared to those identified via mapping studies (such as GWAS). An analogous approach for microbial biodiversity-ecosystem function studies would be to apply artificial ecosystem selection (sensu (56)) on a given function and compare the taxa that change in response to selection with those identified via a comparative approach (such as the one illustrated in our example).

The most common way that loci identified in a mapping study are verified is through manipulative genetics. The identified loci can be knocked out or over-expressed, and the effect on phenotype compared to that predicted from mapping studies. There is no direct analogue for this in microbial biodiversity-ecosystem function studies. In some cases it may be possible to inhibit a particular functional group through the use of a specific antimicrobial or a chemical inhibitor (57), but this is not generally true. It may be possible in some cases to isolate a microorganism of interest in pure culture and add it back to a particular ecosystem, transiently increasing its abundance (roughly analogous to “overexpressing” a gene). Synthetic communities (contrived assemblages of microorganisms) may ultimately be the most powerful way to test hypotheses about microbial biodiversity-ecosystem function relationships, but currently these approaches are limited by the small number of taxa that can be routinely cultured from most environments.

## 4 Conclusion

Microbial biodiversity-ecosystem function research has demonstrated that functional group abundance (measured via genetic markers from environmental DNA) is often not a good predictor of ecosystem function and that microbial diversity metrics on average do not add much power to ecosystem models. A new perspective on how to determine the relationship between microbial communities and ecosystem functions is sorely needed. Organismal biologists have over a hundred years of experience identifying relationships between complex sets of highly-variable data (genotypes or genome sequences) and aggregate functions (organismal phenotypes). We assert that combining the approaches of traditional biodiversity-ecosystem function research with ideas from genotype-phenotype mapping could provide this new perspective. This integration could not only make underutilized approaches such as covariate modeling and artificial selection more available to microbial ecologists, but also provide instructive examples of how best to conceive of microbial biodiversity-ecosystem function questions. If this integration is successful, it is possible that in the not-so-distant future our field will be able to robustly identify taxa, genes, or even molecules that will allow us to accurately predict the response of ecosystems to environmental change. Doing so will not only generate novel hypotheses about how complex microbial communities drive ecosystem function, but also help scientists predict and manage changes to ecosystem functions resulting from human activities.

## Acknowledgments

This project was supported by the National Science Foundation – Dimensions of Biodiversity program (DEB 14422214), the National Science Foundation Graduate Research Fellowship Program (DGE 1255832), and the ARCS Foundation Florence and Mike Nudelman Scholarship. The data re-analyzed here were generated with permission of the Government of Gabon, Centre National de la Recherche Scientifique et Technologique (Permit No AR0035/14/MESRS/CENAREST/CG/CST/CSAR) and the assistance of F. Bivigou, H. Memiaghe, L. Tchignoumba, E. Tobi, and I. Akendengue. The University of Oregon, the Gabon-Oregon Transnational Center on Environment and Development, the Smithsonian Conservation Biology Institute, and Shell Gabon provided financial and logistical support for the original study re-analyzed here. We also thank W. Cresko for insightful discussions on population genetics and H. Tavalire for contributions to the statistical analyses.

## References

1. Hooper DU, Chapin FS, Ewel JJ, Hector A, Inchausti P, Lavorel S, et al. EFFECTS OF BIODIVERSITY ON ECOSYSTEM FUNCTIONING: A CONSENSUS OF CURRENT KNOWLEDGE. Ecological Monographs. 2005 Feb;75(1):3–35, doi:10.1890/04-0922.

2. Schimel JP, Gulledge J. Microbial community structure and global trace gases. Global Change Biology. 1998;4:745–58, doi:10.1046/j.1365-2486.1998.00195.x.

3. Singh BK, Bardgett RD, Smith P, Reay DS. Microorganisms and climate change: terrestrial feedbacks and mitigation options. Nature Reviews Microbiology. 2010 Nov;8(11):779–90, doi:10.1038/nrmicro2439.

4. Rocca JD, Hall EK, Lennon JT, Evans SE, Waldrop MP, Cotner JB, et al. Relationships between protein-encoding gene abundance and corresponding process are commonly assumed yet rarely observed. The ISME Journal. 2015 Aug;9(8):1693–9, doi:10.1038/ismej.2014.252.

5. Graham EB, Knelman JE, Schindlbacher A, Siciliano S, Breulmann M, Yannarell A, et al. Microbes as Engines of Ecosystem Function: When Does Community Structure Enhance Predictions of Ecosystem Processes? Front Microbiol [Internet]. 2016 [cited 2019 Aug 15];7(214), doi:10.3389/fmicb.2016.00214. Available from: https://www.frontiersin.org/articles/10.3389/fmicb.2016.00214/full

6. Reed HE, Martiny JBH. Testing the functional significance of microbial composition in natural communities. FEMS Microbiol Ecol. 2007 Nov 1;62(2):161–70, doi:10.1111/j.1574-6941.2007.00386.x.

7. Glassman SI, Weihe C, Li J, Albright MBN, Looby CI, Martiny AC, et al. Decomposition responses to climate depend on microbial community composition. Proceedings of the National Academy of Sciences. 2018 Nov 20;115(47):11994–9, doi:10.1073/pnas.1811269115.

8. Strickland MS, Lauber C, Fierer N, Bradford MA. Testing the functional significance of microbial community composition. Ecology. 2009;90(2):441–51, doi:10.1890/08-0296.1.

9. Panke-Buisse K, Poole AC, Goodrich JK, Ley RE, Kao-Kniffin J. Selection on soil microbiomes reveals reproducible impacts on plant function. The ISME Journal. 2015 Apr;9(4):980–9, doi:10.1038/ismej.2014.196.

10. Balser TC, Firestone MK. Linking microbial community composition and soil processes in a California annual grassland and mixed-conifer forest. Biogeochemistry. 2005 Apr 1;73(2):395–415, doi:10.1007/s10533-004-0372-y.

11. Cavigelli MA, Robertson GP. The Functional Significance of Denitrifier Community Composition in a Terrestrial Ecosystem. Ecology. 2000 May 1;81(5):1402–14, doi:10.1890/0012-9658(2000)081[1402:TFSODC]2.0.CO;2.

12. Freitag TE, Prosser JI. Correlation of Methane Production and Functional Gene Transcriptional Activity in a Peat Soil. Appl Environ Microbiol. 2009 Nov 1;75(21):6679–87, doi:10.1128/AEM.01021-09.

13. Freitag TE, Toet S, Ineson P, Prosser JI. Links between methane flux and transcriptional activities of methanogens and methane oxidizers in a blanket peat bog. FEMS Microbiol Ecol. 2010 Jul 1;73(1):157–65, doi:10.1111/j.1574-6941.2010.00871.x.

14. Schnyder E, Bodelier PLE, Hartmann M, Henneberger R, Niklaus PA. Positive diversity-functioning relationships in model communities of methanotrophic bacteria. Ecology. 2018 Mar;99(3):714–23, doi:10.1002/ecy.2138.

15. Wagg C, Bender SF, Widmer F, Heijden MGA van der. Soil biodiversity and soil community composition determine ecosystem multifunctionality. PNAS. 2014 Apr 8;111(14):5266–70, doi:10.1073/pnas.1320054111.

16. Maron P-A, Sarr A, Kaisermann A, Lévêque J, Mathieu O, Guigue J, et al. High Microbial Diversity Promotes Soil Ecosystem Functioning. Appl Environ Microbiol. 2018 May 1;84(9):e02738–17, doi:10.1128/AEM.02738-17.

17. Philippot L, Spor A, Hénault C, Bru D, Bizouard F, Jones CM, et al. Loss in microbial diversity affects nitrogen cycling in soil. The ISME Journal. 2013 Aug;7(8):1609–19, doi:10.1038/ismej.2013.34.

18. Hug LA, Baker BJ, Anantharaman K, Brown CT, Probst AJ, Castelle CJ, et al. A new view of the tree of life. Nature Microbiology. 2016 May;1(5):16048, doi:10.1038/nmicrobiol.2016.48.

19. Steen AD, Crits-Christoph A, Carini P, DeAngelis KM, Fierer N, Lloyd KG, et al. High proportions of bacteria and archaea across most biomes remain uncultured. ISME J. 2019 Aug 6;1–5, doi:10.1038/s41396-019-0484-y.

20. Bižić-Ionescu M, Klintzsch T, Ionescu D, Hindiyeh MY, Günthel M, Muro-Pastor AM, et al. Widespread methane formation by Cyanobacteria in aquatic and terrestrial ecosystems. bioRxiv. 2019 Jul 8;398958, doi:10.1101/398958.

21. Lenhart K, Bunge M, Ratering S, Neu TR, Schüttmann I, Greule M, et al. Evidence for methane production by saprotrophic fungi. Nature Communications. 2012 Sep 4;3:1046, doi:10.1038/ncomms2049.

22. Perneger TV. What’s wrong with Bonferroni adjustments. BMJ. 1998 Apr 18;316(7139):1236–8.

23. Rothman KJ. No Adjustments Are Needed for Multiple Comparisons. Epidemiology. 1990;1(1):43–6.

24. Thomas DC, Siemiatycki J, Dewar R, Robins J, Goldberg M, Armstrong BG. THE PROBLEM OF MULTIPLE INFERENCE IN STUDIES DESIGNED TO GENERATE HYPOTHESES. Am J Epidemiol. 1985 Dec 1;122(6):1080–95, doi:10.1093/oxfordjournals.aje.a114189.

25. Benjamini Y, Hochberg Y. Controlling the False Discovery Rate: A Practical and Powerful Approach to Multiple Testing. Journal of the Royal Statistical Society Series B (Methodological). 1995;57(1):289–300.

26. Storey JD. A direct approach to false discovery rates. Journal of the Royal Statistical Society: Series B (Statistical Methodology). 2002 Aug;64(3):479–98, doi:10.1111/1467-9868.00346.

27. Martiny JBH, Bohannan BJM, Brown JH, Colwell RK, Fuhrman JA, Green JL, et al. Microbial biogeography: putting microorganisms on the map. Nature Reviews Microbiology. 2006 Feb;4(2):102–12, doi:10.1038/nrmicro1341.

28. Wright S. Isolation by Distance. Genetics. 1943 Mar 29;28(2):114–38.

29. Berg JJ, Harpak A, Sinnott-Armstrong N, Joergensen AM, Mostafavi H, Field Y, et al. Reduced signal for polygenic adaptation of height in UK Biobank. Nordborg M, McCarthy MI, Nordborg M, Barton NH, Hermisson J, editors. eLife. 2019 Mar 21;8:e39725, doi:10.7554/eLife.39725.

30. Novembre J, Johnson T, Bryc K, Kutalik Z, Boyko AR, Auton A, et al. Genes mirror geography within Europe. Nature. 2008 Nov;456(7218):98–101, doi:10.1038/nature07331.

31. Kang HM, Sul JH, Service SK, Zaitlen NA, Kong S, Freimer NB, et al. Variance component model to account for sample structure in genome-wide association studies. Nature Genetics. 2010 Apr;42(4):348–54, doi:10.1038/ng.548.

32. Price AL, Patterson NJ, Plenge RM, Weinblatt ME, Shadick NA, Reich D. Principal components analysis corrects for stratification in genome-wide association studies. Nature Genetics. 2006 Aug;38(8):904–9, doi:10.1038/ng1847.

33. Meyer KM, Hopple AM, Klein AM, Morris AH, Bridgham S, Bohannan BJM. Community structure – ecosystem function relationships in the Congo Basin methane cycle depend on the physiological scale of function. bioRxiv. 2019 May 17;639989, doi:10.1101/639989.

34. Qin J, Li Y, Cai Z, Li S, Zhu J, Zhang F, et al. A metagenome-wide association study of gut microbiota in type 2 diabetes. Nature. 2012 Sep 26;490(7418):55–60, doi:10.1038/nature11450.

35. Callahan BJ, McMurdie PJ, Holmes SP. Exact sequence variants should replace operational taxonomic units in marker-gene data analysis. The ISME Journal. 2017 Dec;11(12):2639–43, doi:10.1038/ismej.2017.119.

36. McMurdie PJ, Holmes S. phyloseq: An R Package for Reproducible Interactive Analysis and Graphics of Microbiome Census Data. PLOS ONE. 2013 Apr 22;8(4):e61217, doi:10.1371/journal.pone.0061217.

37. R Core Team. R: A language and environment for statistical computing [Internet]. Vienna, Austria: R Foundation for Statistical Computing; 2019. Available from: https://www.R-project.org/

38. Love MI, Huber W, Anders S. Moderated estimation of fold change and dispersion for RNA-seq data with DESeq2. Genome Biology. 2014 Dec 5;15(12):550, doi:10.1186/s13059-014-0550-8.

39. McMurdie PJ, Holmes S. Waste Not, Want Not: Why Rarefying Microbiome Data Is Inadmissible. PLOS Computational Biology. 2014 Apr 3;10(4):e1003531, doi:10.1371/journal.pcbi.1003531.

40. Willis A, Bunge J. Estimating diversity via frequency ratios. Biometrics. 2015;71(4):1042–9, doi:10.1111/biom.12332.

41. Bray JR, Curtis JT. An Ordination of the Upland Forest Communities of Southern Wisconsin. Ecological Monographs. 1957;27(4):325–49, doi:10.2307/1942268.

42. Oksanen J, Blanchet FG, Friendly M, Kindt R, Legendre P, McGlinn D, et al. vegan: Community Ecology Package [Internet]. 2019 [cited 2019 Aug 13]. Available from: https://CRAN.R-project.org/package=vegan

43. Qu L, Guennel T, Marshall SL. Linear Score Tests for Variance Components in Linear Mixed Models and Applications to Genetic Association Studies: Linear Score Tests for Variance Components. Biometrics. 2013 Dec;69(4):883–92, doi:10.1111/biom.12095.

44. Wickham H, Chang W, Henry L, Pedersen TL, Takahashi K, Wilke C, et al. ggplot2: Create Elegant Data Visualisations Using the Grammar of Graphics [Internet]. 2019 [cited 2019 Aug 13]. Available from: https://CRAN.R-project.org/package=ggplot2

45. Belova SE, Pankratov TA, Detkova EN, Kaparullina EN, Dedysh SN. Acidisoma tundrae gen. nov., sp. nov. and Acidisoma sibiricum sp. nov., two acidophilic, psychrotolerant members of the Alphaproteobacteria from acidic northern wetlands. International Journal of Systematic and Evolutionary Microbiology,. 2009;59(9):2283–90, doi:10.1099/ijs.0.009209-0.

46. Chang Y, Land M, Hauser L, Chertkov O, Del Rio TG, Nolan M, et al. Non-contiguous finished genome sequence and contextual data of the filamentous soil bacterium Ktedonobacter racemifer type strain (SOSP1-21T). Stand Genomic Sci. 2011 Oct 1;5(1):97–111, doi:10.4056/sigs.2114901.

47. Domeignoz-Horta LA, DeAngelis KM, Pold G. Draft Genome Sequence of Acidobacteria Group 1 Acidipila sp. Strain EB88, Isolated from Forest Soil. Microbiol Resour Announc. 2019 Jan 3;8(1):e01464–18, doi:10.1128/MRA.01464-18.

48. Fritz I, Strömpl C, Abraham W-R. Phylogenetic relationships of the genera Stella, Labrys and Angulomicrobium within the ‘Alphaproteobacteria’ and description of Angulomicrobium amanitiforme sp. nov. International Journal of Systematic and Evolutionary Microbiology,. 2004;54(3):651–7, doi:10.1099/ijs.0.02746-0.

49. Meyer KM, Memiaghe H, Korte L, Kenfack D, Alonso A, Bohannan BJM. Why do microbes exhibit weak biogeographic patterns? The ISME Journal. 2018 Jun;12(6):1404–13, doi:10.1038/s41396-018-0103-3.

50. Vellend M. Conceptual Synthesis in Community Ecology. The Quarterly Review of Biology. 2010 Jun;85(2):183–206, doi:10.1086/652373.

51. Kerem B, Rommens JM, Buchanan JA, Markiewicz D, Cox TK, Chakravarti A, et al. Identification of the cystic fibrosis gene: genetic analysis. Science. 1989 Sep 8;245(4922):1073–80, doi:10.1126/science.2570460.

52. MacDonald ME, Novelletto A, Lin C, Tagle D, Barnes G, Bates G, et al. The Huntington’s disease candidate region exhibits many different haplotypes. Nat Genet. 1992 May;1(2):99–103, doi:10.1038/ng0592-99.

53. Hindorff LA, Sethupathy P, Junkins HA, Ramos EM, Mehta JP, Collins FS, et al. Potential etiologic and functional implications of genome-wide association loci for human diseases and traits. PNAS. 2009 Jun 9;106(23):9362–7, doi:10.1073/pnas.0903103106.

54. Reich DE, Lander ES. On the allelic spectrum of human disease. Trends in Genetics. 2001 Sep 1;17(9):502–10, doi:10.1016/S0168-9525(01)02410-6.

55. McCalley CK, Woodcroft BJ, Hodgkins SB, Wehr RA, Kim E-H, Mondav R, et al. Methane dynamics regulated by microbial community response to permafrost thaw. Nature. 2014 Oct;514(7523):478–81, doi:10.1038/nature13798.

56. Swenson W, Wilson DS, Elias R. Artificial ecosystem selection. PNAS. 2000 Aug 1;97(16):9110–4, doi:10.1073/pnas.150237597.

57. Maxson T, Mitchell DA. Targeted treatment for bacterial infections: prospects for pathogen-specific antibiotics coupled with rapid diagnostics. Tetrahedron. 2016 Jun 23;72(25):3609–24, doi:10.1016/j.tet.2015.09.069.

